# Local demyelination rekindles oligodendrocyte precursor dynamics and remyelination in old age

**DOI:** 10.64898/2026.02.15.705983

**Authors:** Katharina Eichenseer, Shahrzad Askari, Llazar Shkurti, Thomas Misgeld, Nicolas Snaidero

## Abstract

Cortical myelination is critical for circuit function, plasticity, and long-term stability in the adult brain. With age and disease, the capacity to restore myelin after oligodendrocyte (OL) loss declines, and this failure is thought to reflect intrinsic limitations of OL precursor cells (OPCs) and a loss of permissive cues within the cortical environment. Here, using single-OL ablations and intravital imaging in mice, we show that as cortical remyelination efficiency declines, OPC motility declines, a phenomenon that can be mimicked by blocking CXCL12/CXCR4 signaling. Counter to prevailing notions, however, we find that even in aging, OPCs retain the capacity to generate new OLs and sheath axons, which can be rekindled by a graded demyelinating stimulus, inducing a more juvenile OPC motile state. This reveals an unrecognized remyelination potential in aging and identifies OPC dynamics as a key determinant of cortical remyelination, a property that could be targeted to improve myelin repair.

## Introduction

Oligodendrocytes (OLs) myelinate axons to increase conduction velocity and provide metabolic support, not only in white matter tracts, but also in cortical gray matter (Stadelmann et al., 2019). During primary myelination, OLs derive from precursor cells (OPCs) that dynamically survey their environment (Hughes et al., 2013; Kirby et al., 2006). Additionally, OPCs contribute to ongoing “adaptive” myelination in the adult cortex, supporting cognitive functions such as learning and memory (McKenzie et al., 2014; Steadman et al., 2020). Given these fundamental roles in cortical circuit formation and function, gray matter myelin pathology is now recognized as a major driver of progression in demyelinating diseases such as multiple sclerosis (MS) and hence as a prime therapy target (Albert et al., 2007; Calabrese et al., 2015; Lubetzki et al., 2020). Moreover, myelin deficits have also been described in various neurodegenerative disorders, many of which emerge late in life, including Alzheimer’s disease (Depp et al., 2023). This leads to the important question, whether and how cortical myelination and OPC dynamics can be maintained throughout life.

Cortical myelin stands apart from classical white matter myelin in that it forms complex patterns of internodes along axons, which can exhibit substantial plasticity (Bacmeister et al., 2022; Ford et al., 2015; Hughes et al., 2018; Mitew et al., 2018; Tomassy et al., 2014). These patterns vary between neuronal subtypes (Call & Bergles, 2021; Micheva et al., 2016), indicating a dependence on cell-type-specific signaling and function. The axonal myelination cues that shape these patterns seem to persist into adulthood, as cortical myelin plasticity can continue until mice reach 24 months of age (Hill et al., 2018). Moreover, a specific subset of the cortical myelination patterns can be faithfully reproduced after a bout of demyelination in mature neuropil. Even minimal myelin loss suffices to elicit the targeted replacement of internodes on previously myelinated axons in young adults (Snaidero et al., 2020), a feature conserved across other demyelination paradigms (Chapman et al., 2023; Orthmann-Murphy et al., 2020). These findings establish that OPCs preferentially restore myelin on axons that were strongly myelinated to begin with, while isolated internodes are less efficiently replaced. Despite the clear drive to maintain and restore specific aspects of the cortical myelination pattern, the biological rationale of this robust CNS repair response remains unclear: In the healthy brain, even in aging, cortical OLs appear to undergo only slow turnover, with renewal rates not exceeding 1.5% per month even after mid-age (Tripathi et al., 2017; Young et al., 2013). Thus, OLs – like other glial cells, such as microglia (Askew et al., 2017; Tay et al., 2017) or OPCs (Kirby et al., 2006)– might undergo some level of homeostatic renewal. Still, the turnover rate for OLs is even lower at younger ages (Tripathi et al., 2017) when OL differentiation and myelin production are robust, suggesting that the juvenile potential for precise remyelination is not primarily homeostatic, but rather reserved to enable remodeling and repair. Accordingly, the dynamic behavior of OPCs might serve a sentinel function in an ongoing search for myelination targets, whether newly tagged by circuit plasticity or vacated by demyelinating insults.

Importantly, the dynamic potential and resilience of myelin in young cortex contrasts sharply with the situation in advanced age, when cortical remyelination typically fails. This is true in diseased cortex (Rodriguez et al., 2014) but might also be part of healthy aging (Chapman et al., 2023). Reduced myelin plasticity and repair could contribute to cognitive decline (Gong et al., 2023; Pan et al., 2020; Wang et al., 2020). The underlying reasons are likely multifaceted and include precursor cell-intrinsic factors, such as OPC senescence (Heo et al., 2025; Neumann et al., 2019), as well as extrinsic influences. Here, a leading hypothesis is that the aged neuropil is not permissive to remyelination, with mechanical resistance sensed by OPCs via mechanoreceptors acting as an impediment to proliferation and differentiation (Segel et al., 2019). In this model, however, substantial knowledge gaps remain, despite the biomedical importance of cortical remyelination and its age-related decline. For instance, it is unclear whether OPC dynamics decline in parallel with remyelination potential and whether the two processes are actually linked. Furthermore, it is unclear whether precise remyelination can be rekindled intrinsically in the aged cortex at all. Indeed, many aspects of the ongoing debates related to cortical myelination dynamics are impacted by the variable degree of parenchymal damage involved in the respective demyelination models used to probe the potential for repair, as scarring and local immune responses are age-dependent and can, in turn, influence the efficiency and mechanism with which myelin is restored (Tiwari et al., 2024). For example, the speed and fidelity of remyelination might be influenced by the micromilieu and the level of axonal damage induced by a model. Furthermore, the source of remyelination might vary depending on the number of damaged but surviving OLs, which can serve as an alternative, albeit often inefficient, source of remyelination (Bacmeister et al., 2020; Mezydlo et al., 2023; Neely et al., 2022).

Here, to disentangle such confounders, we use our “minimal” 2-photon-laser-based OL ablation approach across the murine life span to probe remyelination potential in the absence of substantial parenchymal injury, axon loss, or sublethal OL damage (Snaidero et al., 2020). We demonstrate that cortical remyelination efficiency declines substantially even in middle age, impairing the ability to restore cortical myelin architecture even after a minimal lesion caused by the loss of a single OL. We identify an age-related loss in OPC motility as a key cellular determinant of this failure, while reducing OPC motility in young animals via CXCR4/CXCL12 signaling can phenocopy age-related remyelination inefficiency. Conversely, in aged animals, simply by enlarging the minimal lesion threefold, we can rekindle OPC dynamics and their ability to identify and sheath appropriate axonal targets.

## Results

### Cortical remyelination efficiency is impaired at the cellular and internode level in adulthood

We combined single-OL laser ablation (Snaidero et al., 2020) with chronic two-photon imaging in *Plp:GFP(z)* mice across young (2.5-4 months), mid-aged (13.5-17.5 months), and old (22.5-26 months) cohorts to examine age-dependent changes in cortical remyelination (Fig. 1; Extended Data Fig. 1). OLs and their myelin internodes were followed for more than 80 days, capturing acute myelin loss and subsequent remyelination with single-internode resolution (Fig. 1a,b; Extended Data Fig. 1a).

**Figure 1.**
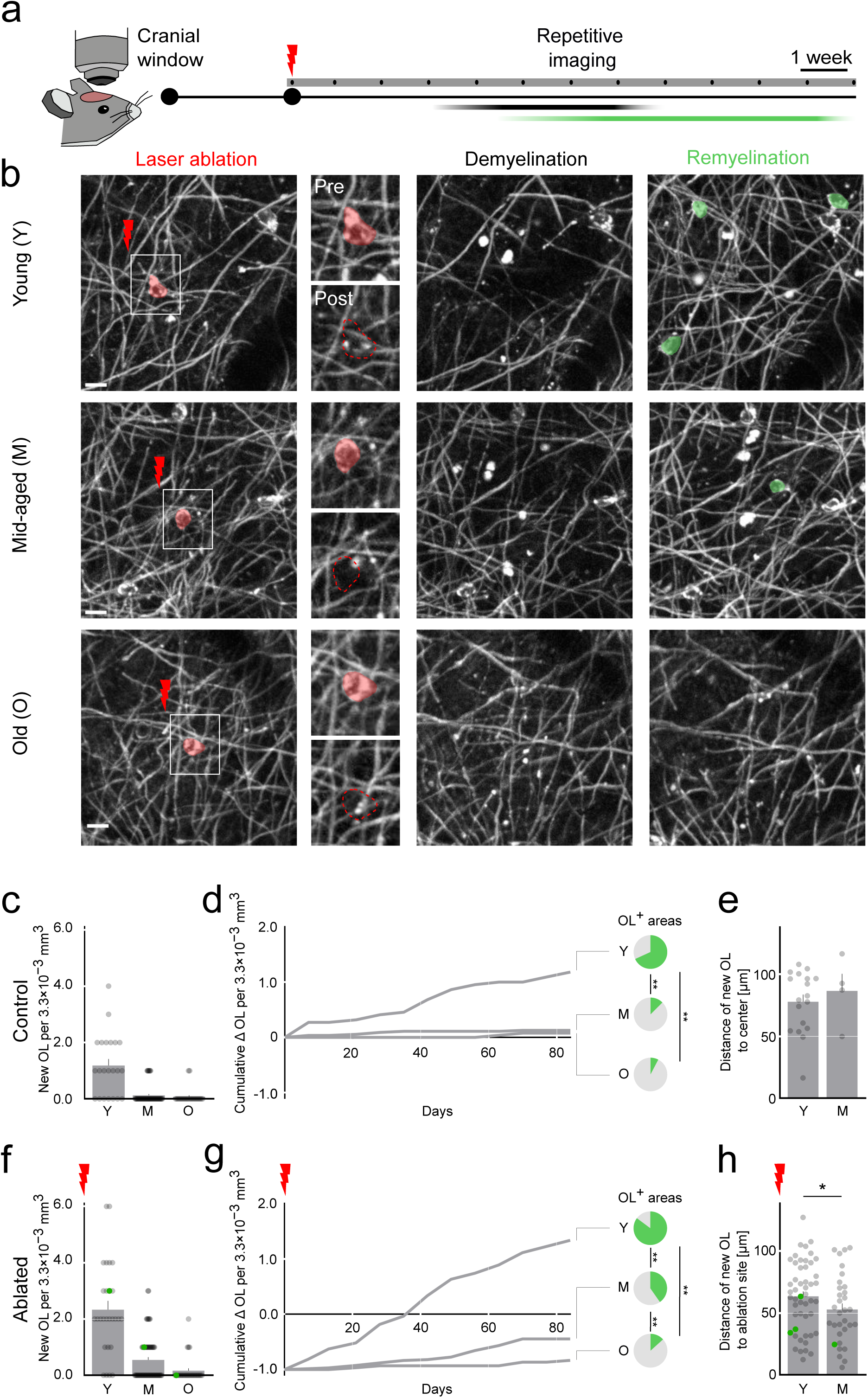
Remyelination efficiency is already impaired in mid-aged animals. **a**, Experimental timeline. Focal laser ablation (red flash) of single oligodendrocytes (OLs) in mice with a chronic cranial window; longitudinal imaging (black dots) performed at least weekly for up to 13 weeks. Periods corresponding to de- and remyelination are indicated by black and green bars, respectively. **b,** In vivo imaging in young (top), mid-aged (middle), and old (bottom) mice. Left, area before single-cell ablation with the target OL highlighted in red; the boxed region is shown at higher magnification in the inset. Insets show the same region before ablation (top) and the position of the ablated OL after ablation (bottom, dashed red outline). Middle, demyelinated region following OL ablation. Right, remyelinated region with newly differentiated OLs shown in green if applicable. Not the full area is shown, maximum-intensity projections, 7.5 µm. **c**, Number of newly differentiated OLs per 3.3×10^−3^ mm^3^ in control regions without laser ablation across age groups (Y, young; M, mid-aged; O, old). **d**, Cumulative number of newly differentiated OLs per 3.3×10^−3^ mm^3^ over time in control regions without laser ablation across age groups. Pie charts show the fraction of “OL+ areas” in control situation (areas with at least one new OL appearing during imaging) in young (top), mid-aged (middle) and old (bottom) animals **e**, Average distance [in µm] of newly differentiated OLs appearing after day 28 from the centered OL in control regions, measured from the position at day 0. **f**, Number of newly differentiated OLs per 3.3×10^−3^ mm^3^ in regions following single-cell laser ablation across age groups. **g**, Cumulative number of newly differentiated OLs per 3.3×10^−3^ mm^3^ over time in regions following single-cell laser ablation. Pie charts show the fraction of “OL+ areas” after OL ablation (areas with at least one new OL appearing during imaging) in young (top), mid-aged (middle) and old (bottom) animals **h**, Average distance [in µm] of newly differentiated OLs appearing after day 28 from the location of the ablated OL at day 0. Bars = mean ± SEM; dots = individual imaging areas (c,g) or individual OLs (f,j). Green colored dots indicate values corresponding to the representative images shown in b. n = young: 22 control areas (18 new OLs post D28) and 27 ablated areas (50 new OLs post D28) from 8 mice; mid aged: 55 control areas (4 new OLs post D28) and 67 ablated areas (31 new OLs post D28) from 18 mice; old: 26 control and 30 ablated areas from 10 mice. Statistics: two-sided Fisher’s exact test and Bonferroni correction for multiple comparison (e,i), unpaired one-tailed t test (f,j). Significance is indicated as P < 0.05 (*) and P < 0.01 (**); higher significance levels are collapsed into **. Exact P values: d, Y vs M, P <0.0001, Y vs O, P <0.0001, M vs O, P = 0.71; e, Y vs M, P = 0.269; g, Y vs M, P <0.0001, Y vs O, P <0.0001, M vs O, P = 0.009; h, Y vs M, P = 0.047 Scale bar, 10 µm (b). See Methods for details.

At the cellular level, OL replacement showed a pronounced age dependence. In young adults, new OL were continuously incorporated under physiological conditions (1.18 ± 0.23 new OLs in our 3.3×10^-3^ mm^3^ sampling volume; Fig. 1c; Extended Data Fig. 1a), and single-cell ablation triggered efficient replacement (2.33 ± 0.31 new OLs; Fig. 1f). In contrast, mid-aged animals exhibited a marked reduction in baseline OL addition (0.13 ± 0.05 new OLs; Fig. 1c, Extended Data Fig. 1b) and ablation-induced replacement (0.55 ± 0.09 new OLs; Fig. 1f, Extended Data Fig. 1c). In old animals, ongoing cortical myelination was similarly low (0.08 ± 0.05 new OLs; Fig. 1c), and remyelination after ablation was rare, occurring in only three of thirty imaged regions (0.17 ± 0.08 new OLs; Fig. 1f).

The time-course of OL incorporation under physiological conditions and after ablation (Fig. 1d,g; Extended Data Fig. 1d) showed a clear age-related decline of repair capacity that paralleled primary myelination: In control areas, young mice showed a steady spontaneous increase in OL numbers (Fig. 1d), while baseline incorporation of new OLs was already markedly reduced in mid-aged animals and virtually absent in old mice (Fig. 1d). Following ablation, young mice displayed a robust regenerative response, with complete replacement being achieved after app. 2.5 months (Fig. 1g). In mid-aged animals, remyelination failed to restore pre-ablation OL levels but was still robust, whereas in old mice incorporation after ablation remained minimal, with no substantial increase over time (Fig. 1g). Consistent with these findings, the proportion of imaging areas in which at least one newly generated OL appeared during the imaging period declined with age (Fig. 1d,g, pie charts, Extended Data Fig. 1b,c). Notably, during the phase of ablation-induced remyelination (i.e. from day 28 after ablation onward), the few OLs generated in mid-aged animals were positioned closer to the ablation site than those in young mice (Y: 64.22 ± 4.01 µm; M: 51.87 ± 4.79 µm; Fig. 1h), whereas no age-dependent difference in OL positioning was observed in control regions (Y: 78.98 ± 5.94 µm; M: 87.86 ± 14.01 µm Fig. 1e). This corroborates the notion that even in young animals, when myelination is still ongoing, OL loss still triggers an additional localized myelination drive that boosts repair – a phenomenon that in mid-aged animals can be studied in isolation, as spontaneous OL generation has ceased.

We next examined whether aging also affected restoration at the level of individual myelin internodes in regions that underwent repair (Fig. 2, Extended Data Fig. 2). Three-dimensional reconstructions of internodes dismantled after ablating an OL (“gone”, Fig. 2a,b) and of internodes that were generated during remyelination (“restored”, Fig. 2a,d), revealed stable OL territories across ages, with no major changes in internode number (Y: 50.33 ± 4.57; M: 51.33 ± 3.00; O: 54.00 ± 4.58; Fig. 2c, top) or average internode length per OL (Y: 65.49 ± 2.34 µm; M: 60.07 ± 2.86 µm; O: 61.52 ± 3.34 µm; Fig. 2c, bottom). Remyelination efficiency ̶ defined as the proportion of demyelinated internodes replaced at their original position ̶ was high in young animals (68.66 ± 4.95%; Fig. 2e, top), but significantly reduced in mid-aged (38.57 ± 4.76%; Fig. 2e, top) and old mice (33.36 ± 4.41%; Fig. 2e, top). When comparing internodes of different fates (permanently lost, restored, or formed de novo on previously unmyelinated axons), restored internodes had the tendency of being longer than lost or de novo ones across all ages (Extended Data Fig. 2a,b), in line with previous reports in young animals (Snaidero et al., 2020). At the same time, there was no age-dependency of the length of internodes, which were later restored (Y: 69.56 ± 1.59 µm; M: 70.99 ± 3.45 µm; O: 72.91 ± 4.71 µm; Fig. 2e, bottom).

**Figure 2.**
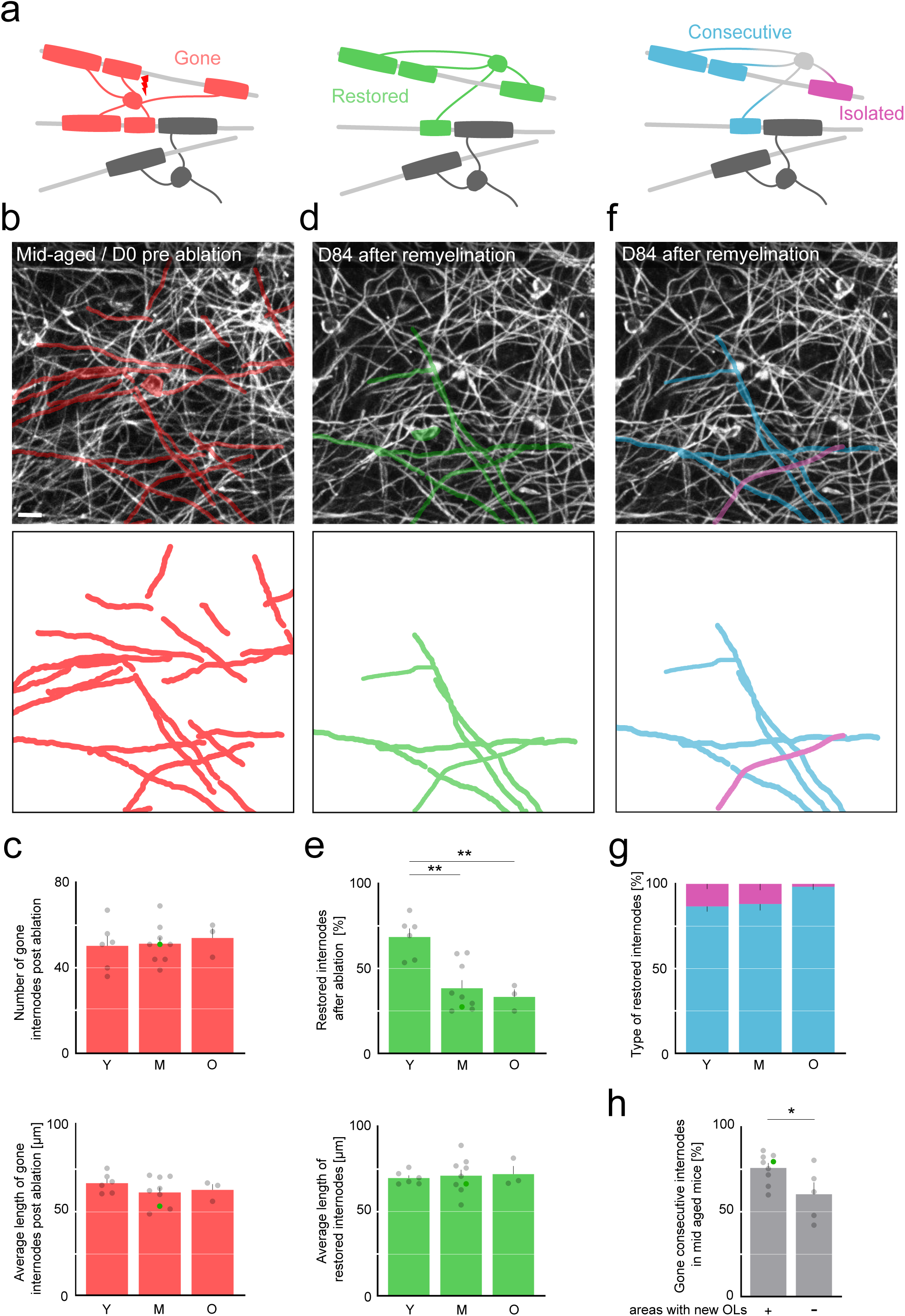
Age-dependent decline in internode restoration and preference for remyelination of consecutive internodes. **a,** Schematic representation of demyelination (left, red), remyelination by a newly generated oligodendrocyte (OL; middle, green), and subsequent classification of restored myelin internodes as consecutive (blue), when flanked by a neighboring internode, or isolated (purple), when occurring alone. Grey OLs and internodes indicate stable OLs; light grey lines represent axons. **b,** Representative in vivo image of an area before ablation in a mid-aged mouse, including tracings internodes (red) that will be demyelinated after single-OL ablation (top). Bottom, representation of only the demyelinated internodes. Maximum-intensity projection, 10 µm. **c,** Number of demyelinated myelin internodes after ablation in young, mid-aged, and old animals (top). Bottom, average length of demyelinated myelin internodes after ablation. **d,** Representative in vivo image of the same area after remyelination in a mid-aged mouse, including tracings of internodes restored after single-OL ablation (top). Bottom, representation of only the restored internodes. Maximum-intensity projection, 10 µm. **e,** Fraction of restored myelin internodes after ablation in young, mid-aged, and old animals (top). Bottom, average length of restored myelin internodes after ablation. **f,** Classification of restored myelin internodes as consecutive (blue) or isolated (purple) (top). Bottom, representation of restored internodes in each category. **g,** Fraction of restored myelin internodes that were consecutive or isolated in young, mid-aged, and old animals. **h,** Fraction of demyelinated consecutive internodes in mid-aged animals, comparing regions that subsequently remyelinated (Areas with new OLs “+”) with regions in which no new OL appeared (Areas with new OLs “-“). Bars = mean ± SEM; dots = individual imaging areas. Colored dots indicate values corresponding to the representative images shown in **b**, **d**, and **f**. *n* = young: 6 areas from 5 mice; mid-aged: 9 areas from 9 mice; old: 3 areas from 2 mice. Statistics: Standard (ordinary) one-way ANOVA with Tukey’s multiple comparisons test (**c**, **e**); unpaired two-tailed *t* test (**h**). Significance is indicated as *P* < 0.05 (*) and *P* < 0.01 (**); higher significance levels are collapsed into **. Exact *P* values: **c** (top), Y vs M, *P* = 0.98; Y vs O, *P* = 0.85; M vs O, *P* = 0.91; **c** (bottom), Y vs M, *P* = 0.37; Y vs O, *P* = 0.73; M vs O, *P* = 0.95; **e** (top), Y vs M, *P* = 0.0013; Y vs O, *P* = 0.037; M vs O, *P* =0.82; **e** (bottom), Y vs M, *P* = 0.95; Y vs O, *P* = 0.92; M vs O, *P* = 0.99; **h**, Areas with new OLs vs Areas without new OLs, *P* = 0.033. Scale bar, 10 µm (**b**). See Methods for details.

To assess whether local axonal context influenced remyelination, internodes were classified as “consecutive” or “isolated” depending on whether they had immediate neighbors (Fig. 2a,f; (Hughes et al., 2018). A comparable fraction of consecutive internodes was demyelinated following OL ablation across age groups (Extended Data Fig. 2e). In all age groups, restoration was strongly biased toward consecutive internodes, whereas isolated internodes were rarely replaced (Y: consecutive 86.68 ± 3.08%, isolated 13.31 ± 3.08%; M: consecutive 88.12 ± 3.79%, isolated 11.88 ± 3.79%; O: consecutive 98.25 ± 1.75%, isolated 1.75 ± 1.75%; Fig. 2g). Thus, although overall remyelination efficiency declined with age, the preferential restoration of consecutive internodes was preserved, as reflected by the higher likelihood of restoring consecutive compared to isolated internodes. In contrast, newly formed internodes generated de novo exhibited an increased fraction of isolated internodes across age groups (Extended Data Fig. 2c,d). When differentiating demyelinated regions by repair outcome, regions that successfully remyelinated in mid-aged animals were enriched in consecutive internodes compared to regions that failed to repair. (Areas with new OLs: 75.92 ± 2.92%; Areas without new OLs: 60.43 ± 6.99%; Fig. 2h).

### OPC dynamics are disrupted early in aging

Having shown that remyelination efficiency declines with age, at both the cellular and internode levels, we next asked whether this deficit originates at the precursor stage. OPCs are known to be dynamic cells, with somata capable of motility and repositioning (Hughes et al., 2013). To assess whether aging alters the in situ behavior of OPCs, we performed longitudinal two-photon imaging in *Plp:GFP* × *Cspg4:dsRed* mice, enabling simultaneous visualization of mature OLs and OPCs (Fig. 3; Extended Data Fig. 3).

**Figure 3.**
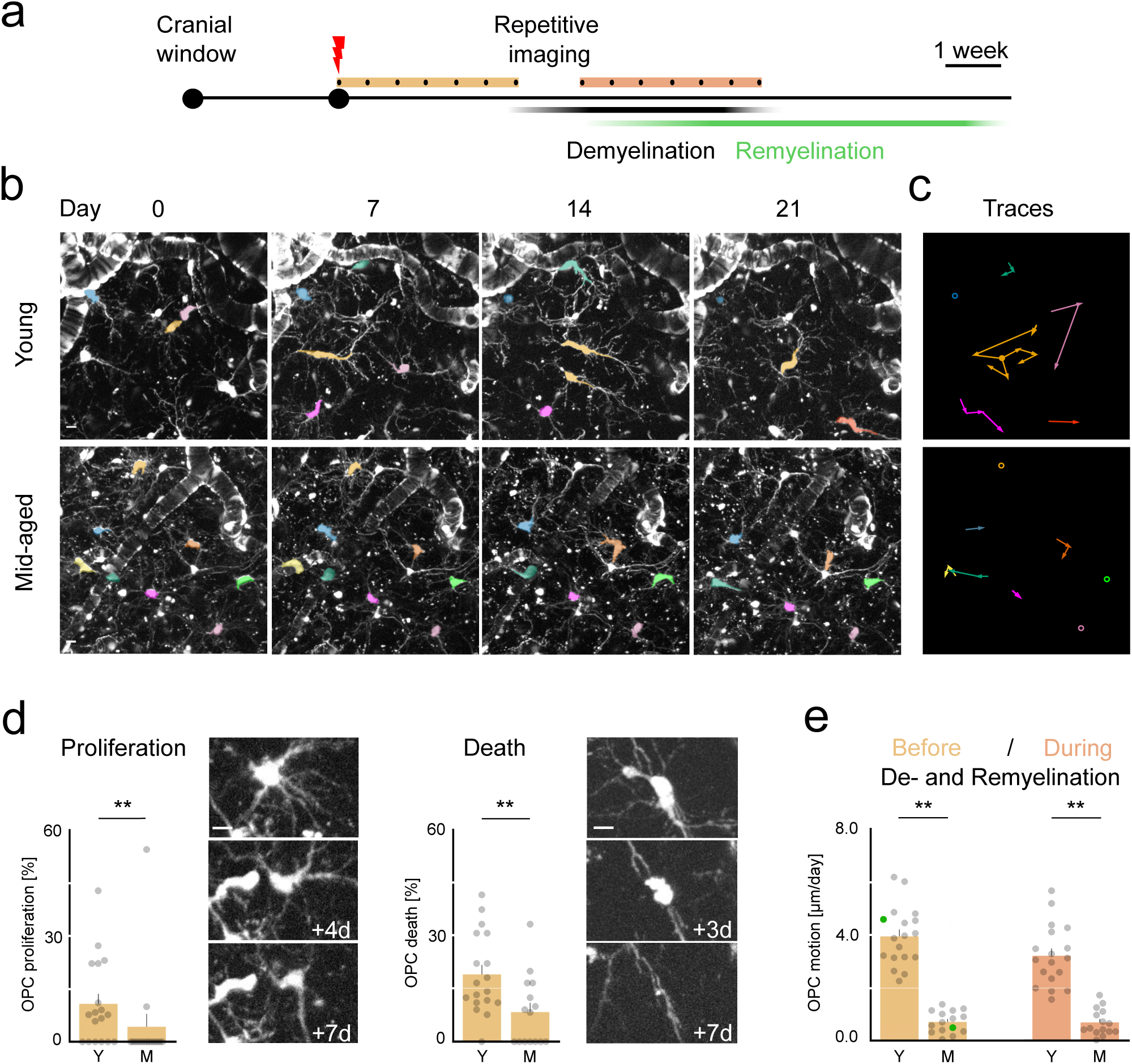
Age-related disruption of OPC dynamics. **a**, Experimental timeline. Focal laser ablation (red flash) of single oligodendrocytes (OLs) in mice with a chronic cranial window. Longitudinal imaging (black dots) used for OPC quantification was performed over the first 3 weeks post-ablation (light orange bar) and during the subsequent de-/remyelination phase (dark orange bar), with imaging 2-3 times per week. Periods corresponding to de- and remyelination are indicated by black and green bars, respectively. **b**, In vivo longitudinal two-photon imaging of OPCs in young (top) and mid-aged (bottom) mice. OPC somata present for at least two consecutive time points were analyzed and pseudo-colored consistently across sessions. Representative images are shown. Maximum-intensity projections, 40 µm. **c**, Trajectories of OPC soma movement derived from the representative images in b. Arrowheads indicate the direction of movement between consecutive time points; open circles indicate stable OPCs, and filled circles indicate proliferation events. **d**, Rates of OPC proliferation (left) and cell death (right) during the first 3 weeks after single-OL ablation. Representative images for 1 week of imaging of young animals are shown to the right of the graphs. Maximum-intensity projections, 37.5 µm (proliferation) and 35 µm (cell death). **e**, Average daily OPC soma displacement in young compared to mid-aged mice before (light orange) and during (dark orange) the de-/remyelination phase. Bars = mean ± SEM; dots = individual imaging areas. Colored dots indicate values corresponding to the representative images shown in b. n = young: 18 ablated areas from 5 mice; mid-aged: 15 ablated areas from 6 mice. Statistics: two-tailed Mann-Whitney U test (d left); non-paired two tailed t-test (d right); Welch’s one-way ANOVA followed by Dunnett’s T3 multiple-comparisons test (e). Significance is indicated as P < 0.05 (*) and P < 0.01 (**); higher significance levels are collapsed into **. Exact P values: d (left), Y vs M, P = 0.007; d (right), Y vs M, P = 0.008; e, Y (pre) vs M (pre), P < 0.0001; Y (pre) vs Y (post), P = 0.30; Y (post) vs M (post), P < 0.0001; M (pre) vs M (post), P > 0.99. All post hoc tests were performed, but only comparisons relevant to the experimental question are shown. Scale bar, 10 µm (b). See Methods for details.

Individual OPCs were followed over three weeks with 2-3 imaging sessions per week, and the soma displacement over time was quantified to assess age-dependent changes in OPC dynamics (Fig. 3a-c). In young cortex, OPC somata displayed pronounced motility in both control and ablated areas across imaging sessions (Fig. 3b, c, top panels; Extended Data Fig. 3a, b, top). This dynamic behavior contrasted sharply with the near standstill observed in mid-aged animals, where soma displacement was reduced more than fivefold (Fig. 3b, c, bottom; Extended Data Fig. 3a, b, bottom). This reduction was consistent and persisted throughout the imaging period (Fig. 3e; Extended Data Fig. 3c). OPC behavior was not significantly altered by laser ablation, local demyelination, or subsequent remyelination after the loss of a single OL, as soma displacement remained comparable between early (day 0-21, after laser ablation) and late (day 28-49, during remyelination) phases and was similarly stable in non-ablated regions (Y, D0-21: 3.95 ± 0.26 µm/day; Y, D28-49: 3.216 ± 0.27 µm/day; M, D0-21: 0.72 ± 0.10 µm/day; M, D28-49: 0.69 ± 0.13 µm/day; Fig. 3e; Extended Data Fig. 3d, e). Despite these substantial changes in dynamic reach, OPC density remained constant across ages (Extended Data Fig. 3f), suggesting that the ability of OPCs to scan the surrounding tissue declines.

Given the pronounced reduction in OPC motility, we next examined whether their subsequent lineage behavior was also altered. OPC proliferation was significantly reduced in mid-aged cortex compared to young animals (Y: 10.78 ± 2.85%; M: 4.30 ± 3.65%; Fig. 3d, left), consistent with previous reports of declining OPC cycling with age (Heo et al., 2025; Segel et al., 2019). In contrast, OPC survival was increased, with fewer cell deaths observed in mid-aged than in young mice (Y: 19.12 ± 2.70%; M: 8.44 ± 2.60%; Fig. 3d, right). Thus, although OPC density remained stable, aging shifted OPC behavior toward a more static state characterized by reduced motility, diminished proliferation, and decreased turnover.

Together, these findings show that OPC dynamics become markedly less active already in midlife, with reduced soma motility and lineage turnover despite preserved cell density.

### Reducing OPC motility impairs remyelination

Given that OPC motility declined sharply in mid-aged cortex, we next asked whether reducing their movement in young animals would be sufficient to impair remyelination. To test this, *Plp:GFP* × *Cspg4:dsRed* mice underwent longitudinal two-photon imaging following single-cell ablation while receiving a pharmacological CXCR4 antagonist (Fig. 4; Extended Data Fig. 4), an intervention that has previously been shown to affect OPC motility in vitro (Dziembowska et al., 2005).

**Figure 4.**
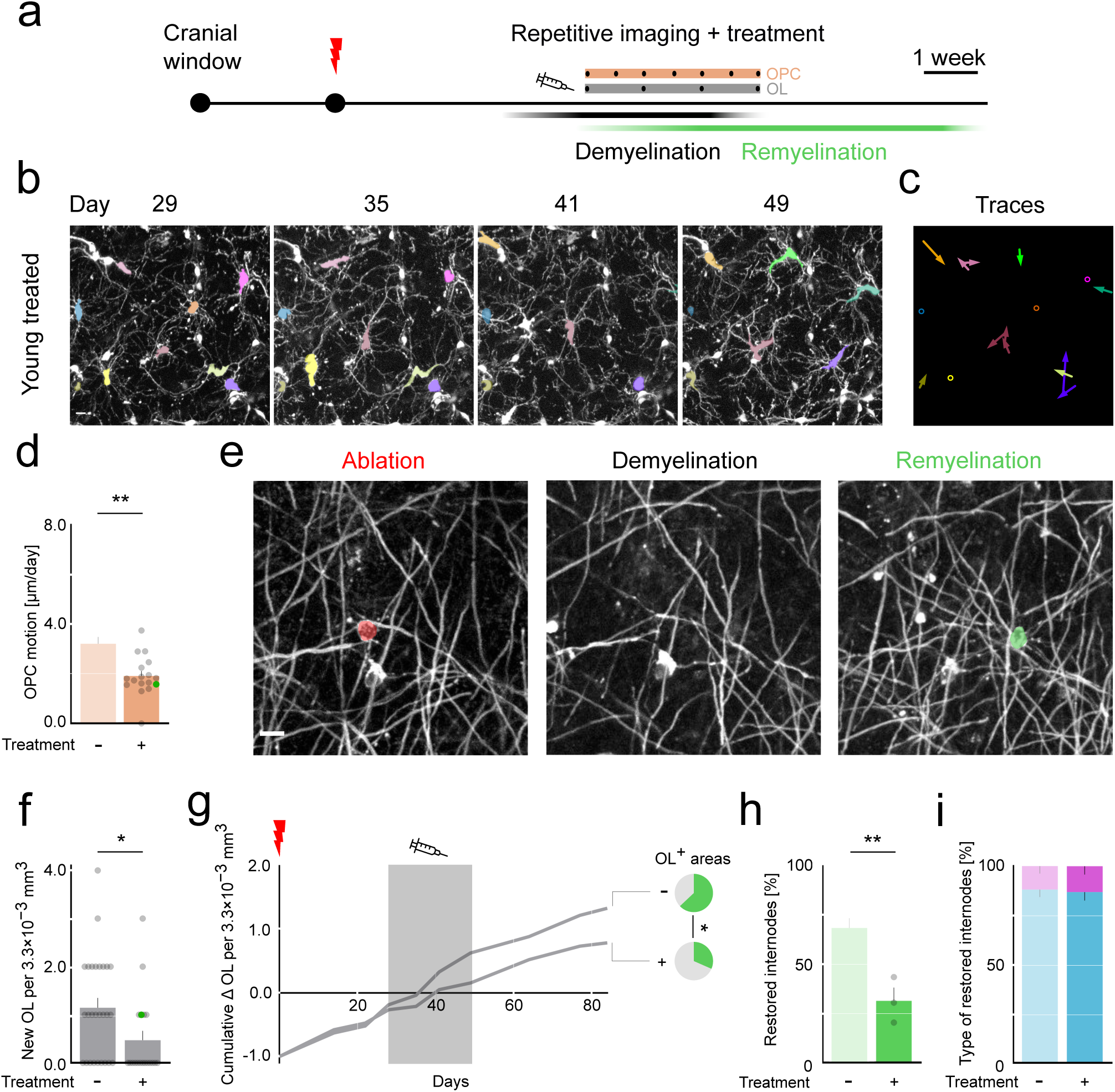
Pharmacological inhibition of CXCR4 reduces OPC motility and impairs remyelination. **a**, Experimental timeline. Focal laser ablation of single oligodendrocytes (OLs) at day 0 in *Plp:GFP* × *Cspg4:dsRed* mice with a chronic cranial window. Analysis was performed over a 3-week period during CXCR4-antagonist treatment, with imaging 2–3 times per week. **b**, In vivo longitudinal two-photon imaging of OPCs in a young treated mouse. OPC somata present for at least two consecutive time points were analyzed and pseudo-colored consistently across sessions. Representative images are shown. Maximum-intensity projections, 40 µm. **c**, Trajectories of OPC soma movement derived from the representative images in b. Arrowheads indicate the direction of movement between consecutive time points; open circles indicate stable OPCs. **d**, Average daily OPC soma displacement during CXCR4-antagonist treatment (control untreated is replotted from Fig. 3e). **e**, In vivo imaging in young treated mouse. Left, area before single-cell ablation with the target OL highlighted in red. Middle, demyelinated region following OL ablation. Right, remyelinated region with newly differentiated OL highlighted in green. Not the full area is shown, maximum-intensity projections, 7.5 µm. **f**, Density of newly differentiated OLs per 3.3×10^−3^ mm^3^ during the CXCR4-antagonist treatment window compared to untreated animals in ablated regions. **g**, Cumulative number of newly differentiated OLs per area over time in untreated (-) (replotted from Fig. 1g) versus treated (+) animals; the treatment window is indicated in grey. Pie charts show the fraction of “OL+ areas” after OL ablation (areas with at least one new OL) considering the time during the treatment (bottom) or comparable time window in control animals (top). **h**, Restoration efficiency (fraction of demyelinated internodes replaced by new OLs), in CXCR4-antagonist treated compared to untreated (replot from Fig. 2e) animals. **i**, Fraction of restored myelin internodes classified as consecutive (blue) or isolated (purple) after ablation in CXCR4-antagonist treated animals and untreated (replot from Fig. 2g) animals. Bars = mean ± SEM; dots = individual imaging areas. Colored dots indicate values corresponding to the representative images shown in d and f. Replotted values from Figs. 1 and 2 are represented without single values and greyed out. n = treated: 17 areas from 6 mice (d); treated: 19 areas from 6 mice and untreated: 27 areas from 8 mice (f,g); treated: 3 areas from 3 mice (h,i). Statistics: non-paired two-tailed t-test (d,f,h), two-sided chi-square test (g, pie charts). Significance is indicated as P < 0.05 (*) and P < 0.01 (**); higher significance levels are collapsed into **. Exact P values: d, – vs +, P < 0.001; f, – vs +, P = 0.017, g, – vs +, P = 0.036; h, – vs +, P = 0.003. Scale bar, 10 µm (b,e). See Methods for details.

Indeed, in vivo tracking and motion reconstructions revealed that CXCR4 inhibition markedly decreased daily soma displacement compared to untreated controls (untreated: 3.22 ± 0.27 µm/day; treated: 1.92 ± 0.20 µm/day; Fig. 4a-d; Extended Data Fig. 4a). This reduction was also evident when comparing OPC motility before and during drug administration in the same animals, confirming a consistent decline independently of laser ablation and OPC number (Extended Data Fig. 4b,c). In parallel, a reduction in newly generated OLs was detected (untreated: 0.35 ± 0.06 OL/mm³; treated: 0.14 ± 0.06 OL/mm³; Fig. 4e,f; Extended Data Fig. 4d). Cumulatively, OL incorporation over time was markedly reduced starting with the CXCR4 inhibition (Fig. 4g). Consistent with this, the fraction of imaging areas in which at least one newly generated OL appeared was significantly reduced under CXCR4 blockade, reflecting a global decrease in OL-generating activity (Fig. 4g, pie charts). At the internode level, while the number and characteristics of demyelinated and newly generated myelin were unchanged per OL (Extended Data Fig. 4g), restoration efficiency was diminished under CXCR4 blockade (untreated: 68.66 ± 4.95%; treated: 31.57 ± 6.71%; Fig. 4h), mirroring the reduction seen in mid-aged animals (cf. Fig. 2e, top). Notably, the preference for consecutive internode restoration remained unchanged, indicating that this organizational rule of remyelination is maintained even when OPC motion is restricted (treated: consecutive 86.02 ± 4.46%, isolated 13.98 ± 4.46%; Fig. 4i; Extended Data Fig. 4h, cf. Fig. 2g).

Together, these results demonstrate that OPC motility is a prerequisite for efficient cortical remyelination. Pharmacological inhibition of CXCR4 in young mice reproduces the cellular-and internode level deficits observed with aging, demonstrating that impaired OPC dynamics can contribute to remyelination failure.

### Local extent of demyelination modulates OPC motility and remyelination

Finally, we examined whether the mid-aged cortex retains the capacity to mount a regenerative response following more extensive demyelination. To this end, we compared the response to ablation of a single OL with that following ablation of three neighboring OLs (Fig. 5a,b; Extended Data Fig. 5). As expected, multiple ablations generated a substantially larger demyelinated area, with a proportional (≍3-fold) increase in lost internodes than after single-OL ablation (single: 51.33 ± 2.98; multi: 149.33 ± 13.25; Fig. 5c). At the cellular level, differentiation (multi: 0.398 ± 0.11 OL/mm³; Fig. 5d) and integration of new OLs increased more steeply following multiple ablations (Fig. 5d). Notably, even very old animals showed a rekindling of remyelination activity, albeit to a weaker extent on a delayed time axis (Extended Data Fig. 5a,b). Together, these data indicate that, despite an overall age-related decline in remyelination capacity, the mid-aged cortical environment retains the ability to reinitiate OL generation when demyelination is sufficiently extensive, in the absence of other changes to the microenvironment or metabolic stress to OPCs, as is likely in most other demyelination models.

**Figure 5.**
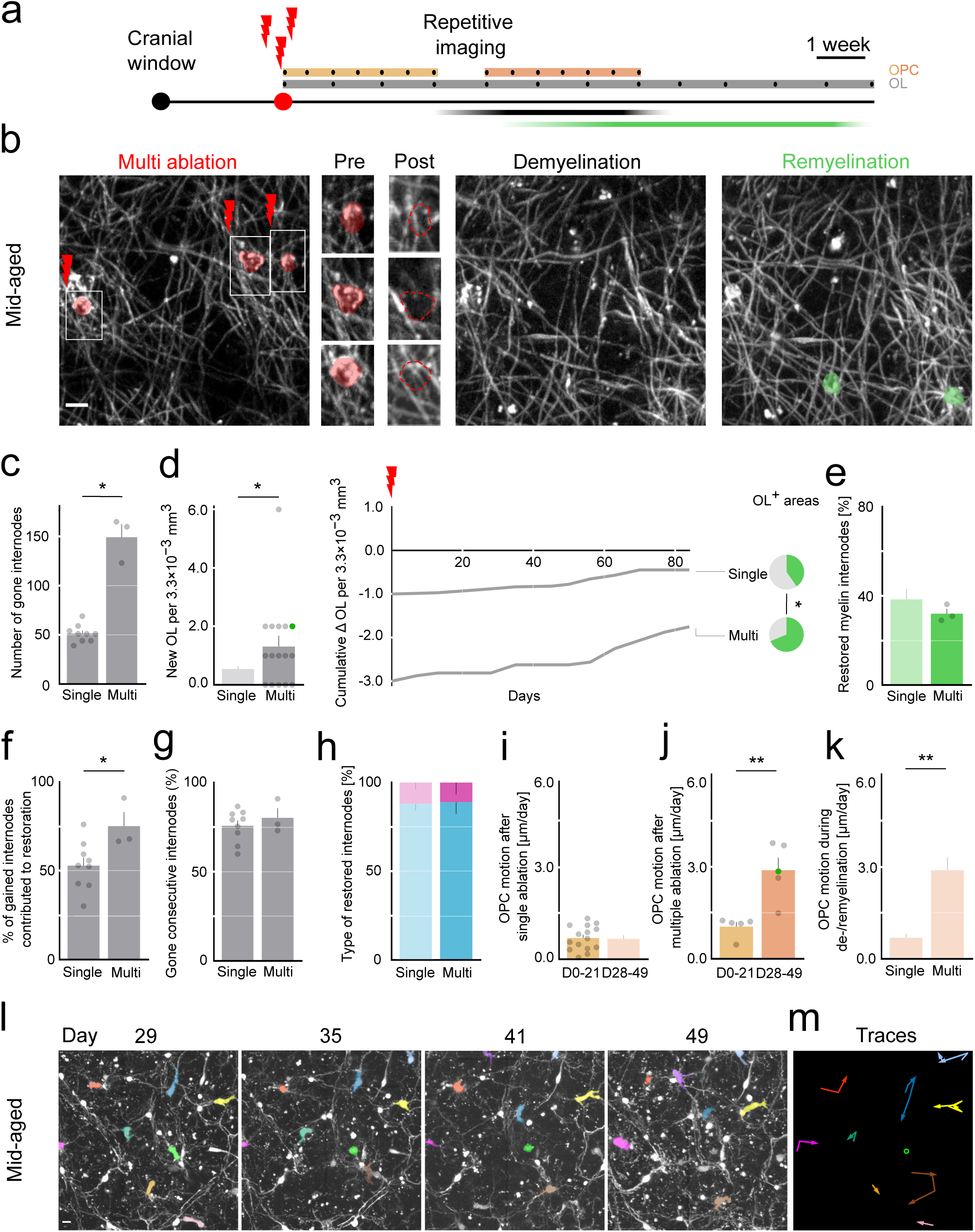
Local extent of demyelination modulates OPC motility and remyelination. **a**, Experimental timeline. Focal ablation of three oligodendrocytes (OLs) at day 0 in *Plp:GFP* × *Cspg4:dsRed* mice with a chronic cranial window. OPC dynamics were analyzed over two 3-week periods: immediately after ablation (light orange) and later during the de-/remyelination phase (dark orange), with imaging performed 2-3 times per week (black dots). OLs were analyzed across the entire imaging period (grey), with weekly imaging time points (black dots). **b**, In vivo imaging in mid-aged mice. Left, area before multiple-cell ablation with the target OLs highlighted in red; boxed regions are shown at higher magnification in the insets. Insets show the same region before ablation (left) and the position of the ablated OLs after ablation (right, dashed red outlines). Middle, demyelinated region following OL ablation. Right, remyelinated region with two newly differentiated OLs shown in green. Maximum-intensity projections, 17.5 µm. **c**, Number of demyelinated myelin internodes after single-cell or multiple-cell ablation in mid-aged animals. **d**, Number of newly differentiated OLs per 3.3×10^−3^ mm^3^ in regions following single-cell (replot from Fig. 1f) and multiple-cell laser ablation. Cumulative number of newly differentiated OLs per 3.3×10^−3^ mm^3^ over time in regions following single-cell (replot from Fig. 1g) and multiple-cell laser ablation. Pie charts show the fraction of “OL+ areas” in single ablated (top, replot from Fig. 1g) and multiple ablated (bottom) animals. **e**, Fraction of restored myelin internodes after single-cell (replot from Fig. 2e) or multiple-cell ablation in mid-aged animals. **f**, Fraction of gained internodes that contributed to remyelination after single-cell or multiple-cell ablation in mid-aged animals. **g**, Fraction of gone internodes, that were consecutive after single-cell or multiple-cell ablation in mid-aged animals. **h**, Fraction of restored myelin internodes that were consecutive or isolated after single-cell (replot from Fig. 2g) or multiple-cell ablation in mid-aged animals. **i**, Average daily OPC soma displacement after single-cell ablation in mid-aged animals (replot from Fig. 3e) right after ablation (D0-21) vs during de-/remyelination phase (D28-49). **j,** Average daily OPC soma displacement after multiple-cell ablation in mid-aged animals right after ablation (D0-21) vs during de-/remyelination phase (D28-49). **k,** Average daily OPC soma displacement over three weeks during the de-/remyelination phase (D28-49), comparing single- and multiple-ablated regions. (Results are replotted from Fig. 5j and Fig. 3e). **l,** In vivo longitudinal two-photon imaging of OPCs in mid-aged multiple-cell ablated mice. OPC somata present for at least two consecutive time points were analyzed and pseudo-colored consistently across sessions. Representative images are shown. Maximum-intensity projections, 40 µm. **m**, Trajectories of OPC soma movement derived from the representative images in k. Arrowheads indicate the direction of movement between consecutive time points; open circles indicate stable OPCs. Bars = mean ± SEM; dots = individual imaging areas. Colored dots indicate values corresponding to the representative images shown in b and k. Replotted values from Fig. 1, 2 and 3 are represented without single values and greyed out; n (OL counts) = mid-aged: 67 single-ablated areas from 19 mice and 16 multiple-ablated areas from 6 mice (d); n (myelin tracing) = 9 single-ablated areas from 9 mice and 3 multiple-ablated areas from 3 mice (e-h); n (OPC motion) = 15 single-ablated areas from 6 mice and 5 multiple-ablated areas from 3 mice (i-k). Statistics: Welch’s two-tailed t test (c,f,g,j); two-tailed Mann-Whitney U test (d left,e,i,k); two-sided chi-square test (d, pie-charts). Significance is indicated as P < 0.05 (*) and P < 0.01 (**); higher significance levels are collapsed into **. Exact P values: c, single vs multiple, P = 0.014; d (left), single vs multiple, P = 0.014; d (pie-charts), single vs multiple, P = 0.04; e, single vs multiple, P > 0.99; f, single vs multiple, P = 0.035; g, single vs multiple, P = 0.53; i, D0-21 vs D28-49, P = 0.84; j, D0-21 vs D28-49, P = 0.009; k, single vs multiple, P < 0.0001. Scale bar, 10 µm (b,k). See Methods for details.

At the internode level, restoration efficiency was not detectably changed following single and multiple ablations (multi: 38.27 ± 4.75%; Fig. 5e). Because the absolute number of lost internodes was substantially higher after multiple ablations (Fig. 5c), maintaining a similar efficiency resulted in a markedly greater number of restored internodes generated by newly differentiated OLs (multi: 75.23 ± 7.93%; Fig. 5f). The length of the internodes and proportion of consecutive internodes lost and newly gained was similar after single and multiple ablations (single: 75.92 ± 2.92%; multi: 80.27 ± 5.45%; Fig. 5g; Extended Data Fig. 5c,d). Moreover, the distribution of restored internode types was unchanged, with restoration remaining strongly biased toward consecutive internodes in both conditions (single: 88.12 ± 3.79% consecutive, 11.88 ± 3.79% isolated; multi: 88.99 ± 6.75% consecutive, 11.01 ± 6.75% isolated; Fig. 5h, Extended Data Fig. 5e).

We next assessed whether this enhanced repair response was accompanied by changes in OPC behavior. Tracking of individual OPC somata revealed that motility did not differ between early and later phases following single ablation (D0-21: 0.722 ± 0.11 µm/day; D28-49: 0.69 ± 0.13 µm/day; Fig. 5i). By contrast, after multiple ablations, OPC soma displacement increased markedly during the subsequent repair phase, while remaining unchanged immediately after ablation (multiple: pre: 1.05 ± 0.15 µm/day; post: 2.88 ± 0.41 µm/day; Fig. 5j). Quantification revealed an approximately fivefold increase in average daily soma displacement under multiple-ablation conditions (multiple: 2.88 ± 0.41 µm/day; Fig. 5k,l,m), indicating that extensive demyelination can re-engage OPC motility even in mid-aged cortex.

Together, these findings demonstrate that increasing the extent of demyelination reactivates OPC motility and rekindles myelin repair.

## Discussion

Using laser ablation of myelinating cells, the present study reveals two key insights. First, even in the aged CNS, precise, quantitative remyelination remains possible, provided a suitable demyelinating stimulus occurs in the absence of general neuropil damage (Fig. 1, 2, 5). Second, the degree to which myelin can be repaired depends on the intrinsic motility of OPCs, which declines with age (and in response to blocking CXCR4 signaling), but returns in response to a sufficient demyelination stimulus (Fig. 3, 4, 5). Together, these findings position OPC dynamics as a central, age-sensitive determinant of cortical remyelination, linking cellular behavior to the tissue’s capacity to respond to specific damage with an appropriate and effective repair response.

Cortical remyelination, which is key for axonal health and cognitive function (Simons & Nave, 2016), has long been considered severely compromised in ageing (Bonetto et al., 2021). Studies using optochemical ablation of individual OLs (Chapman et al., 2023), but also large inflammatory or toxic demyelination models (Neumann et al., 2019) report little effective repair in old animals, supporting the view that intrinsic OPC decline and a less permissive environment limit regeneration (Mironova et al., 2026; Segel et al., 2019). Yet these approaches leave open whether ageing abolishes the remyelination program itself or primarily impairs its reactivation under certain pathological conditions.

Our data argue against a complete loss of competence. While even minimal and “scar-free” focal demyelination reveals a pronounced age-dependent decline in OL replacement and internode restoration, key aspects of demyelination-induced remyelination are preserved even in advanced age. For instance, internode replacement retains spatial specificity, preferentially restoring previously myelinated axons. Thus, while remyelination efficiency declines early quantitatively, the qualitative capacity for targeted repair persists once the process is successfully initiated by an appropriately strong and specific demyelination stimulus.

Three aspects of these results are particularly noteworthy: (1) The age at which damage-induced remyelination declines coincides with the completion of primary cortical myelination in the cortex. (Fig. 1d,g, and (Hughes et al., 2018). This suggests that the age-dependent remyelination capacity of cortical myelin may reflect not only a repair program, but rather the continued availability of a developmentally plastic state characterized by a high number of myelination-competent axonal targets. During early adulthood, ongoing circuit refinement and adaptive myelination expand this pool of available targets, thereby increasing the probability that newly generated OLs can efficiently locate and remyelinate appropriate axons (Bonetto et al., 2021). As cortical circuits stabilize with age and adaptive myelination slows down (Hughes et al., 2018), the availability of myelination-competent targets is progressively reduced, imposing an additional constraint on remyelination efficiency that becomes apparent even when OL differentiation can still occur. (2) Despite the failure of OPC recruitment and differentiation starting in mid-age, the surrounding (and in our model, undamaged) OLs did not mount a repair response (Fig. 1b; cf. (Mezydlo et al., 2023) and remyelination essentially paralleled OL replacement, refuting the notion that in older ages, non-canonical modes of OL-mediated remyelination could dominate (Franklin et al., 2021). (3) Our result that single-cell OL ablations in animals of very advanced age, if at all, only elicit a minimal remyelination response (Fig. 1f,g) chimes with recent optochemical (2Phatal) ablation studies that reported a complete absence of OL differentiation (Chapman et al., 2023). This prior result, together with the absence of substantial repair in large-scale demyelination models in aging rodents, was interpreted to suggest that in old age, OPCs are generally unable to repair myelin loss (Neumann et al., 2019). In contrast, our multi-cell ablations demonstrate that remyelination failure in ageing animals is not absolute but depends on the scale of demyelination, with failure at both extremes: single-cell and global lesions. We interpret this non-linear dependence as indicative of two mechanisms: Lesions that are too small do not elicit a sufficiently strong remyelination signal, because the surrounding tissue response that in younger animals supports remyelination is muted (e.g. by microglia, (Lloyd et al., 2019) or because an insufficient number of myelination-supportive axonal targets is denuded in a neuropil where the density of such targets is physiologically reduced due to progressive myelin addition (Hill et al., 2018). In contrast, in large demyelinating lesions, additional age-related factors such as smoldering inflammation and glial activation (Hammond et al., 2019), extracellular matrix remodeling (Mironova et al., 2026; Segel et al., 2019), or axonal loss (Sim et al., 2002) impose an extrinsic constraint on OPC dynamics and differentiation, thereby suppressing the repair response that larger lesions can elicit even in the aged cortex.

Taken together across the lifespan, our data reveal that while remyelination certainly becomes less efficient in old age, this reduced repair potential is less an abrupt “failure” after a tipping point such as a loss of the intrinsic differentiation capacity of OPCs (Mironova et al., 2026; Neumann et al., 2019) or a fundamental inability of the cortical milieu to support repair (Mironova et al., 2026; Segel et al., 2019). Rather, the persistent capacity for remyelination seems progressively masked as context-dependent constraints on OPC engagement grow and target availability declines. Recent work has proposed that age-related remyelination failure primarily reflects altered spontaneous OL lineage progression, with repair outcomes largely determined by cell-autonomous differentiation programs rather than spatially instructive cues in the tissue environment (Mironova et al., 2026). Our findings are compatible with this model but indicate that altered lineage progression does not fully account for remyelination outcomes following focal myelin loss. In particular, the graded increase in repair probability with increasing focal demyelination argues against a purely stochastic deployment of differentiation events and instead supports the existence of spatial constraints that bias where and when differentiation is engaged. Thus, while intrinsic lineage programs shape OPCs’ baseline propensity to differentiate, our data demonstrate that lesion context remains a critical determinant of spatially targeted remyelination even in advanced age.

Middle age appears to be a critical time of transition, when the rodent cortex ceases to spontaneously myelinate, probably because most constitutive myelination targets have been sheathed and new ones appear at a reduced rate due to diminished circuit plasticity (Burke & Barnes, 2006; McKenzie et al., 2014). Our data reveal that this progressive loss of OPC remyelination efficiency corresponds to a decline in cell body motility, a unique feature of OPCs compared to all other resident cells of the mature CNS outside neurogenic niches (Hughes et al., 2013). Starting at one year of age, murine OPCs show a pronounced reduction of these dynamics, even under physiological conditions. Importantly, impaired OPC dynamics are not merely a correlate of ageing but directly limit remyelination efficiency and can phenocopy key features of age-associated remyelination failure. In young cortex, attenuation of OPC motility via manipulation of the CXCR4/CXCL12 signaling pathway (Dziembowska et al., 2005). is sufficient to markedly reduce remyelination following focal demyelination. This not only reveals the role of OPC dynamics in remyelination but also shows the potential to modulate CXCR4/CXCL12 signaling as one example of a broader class of pathways capable of modulating OPC motility and cortical remyelination. Consistent with this interpretation, we further show that clustered multi-cell OL ablation in advanced age re-engages OPC motility alongside remyelination, indicating that dynamic constraints on OPC behavior remain reversible under appropriate lesion conditions.

Together, these findings suggest that age-related remyelination failure reflects a progressive suppression of OPC dynamics rather than an irreversible loss of repair competence. Crucially, middle age emerges as a window in which declining OPC motility already limits remyelination efficiency yet remains amenable to intervention. Targeting pathways that regulate OPC dynamics, such as CXCR4/CXCL12 signaling, therefore represents a strategy to preserve cortical myelin integrity before repair failure becomes entrenched, and may also be leveraged to unlock residual repair capacity in the aged or diseased brain, especially when combined with approaches that also modify secondary tissue damage (Segel et al., 2019) and boost OPC differentiation (Neumann et al., 2019)

## Materials and methods

### Transgenic animals

Experiments were performed using transgenic mice expressing GFP under the proteolipid protein (PLP) promoter on a C57BL/6 background (here referred to as *Plp:GFP* mice, (Spassky et al., 2001). Transgenic mice expressing dsRed.T1 under the control of the Cspg4 promoter (Jackson Laboratory, #008241; *Cspg4:dsRed*) were also used and maintained in-house. Double-transgenic mice carrying both transgenes (*Plp:GFP × Cspg4:dsRed*) were generated in our animal facilities.

Both male and female mice ranging from 2.5 to 26 months of age were included. Age groups were defined based on the age at the start of imaging as follows: young (2.5-4 months), mid-aged (13.5-17.5 months), and old (22.5-26 months). Animals were housed under controlled conditions (22 ± 2 °C, 50 ± 15% humidity, 12-h light/dark cycle). All procedures complied with German animal welfare regulations (TierSchG) and were approved by the local authorities (Regierung von Oberbayern).

### Cranial window surgery

To enable longitudinal in vivo imaging of OL lineage cells and, a cranial window was implanted above the somatosensory cortex as described previously (Holtmaat et al., 2009; Jafari et al., 2021). Briefly, mice were anesthetized by intraperitoneal injection of medetomidine (0.5 mg/kg), midazolam (5 mg/kg), and fentanyl (0.05 mg/kg). A craniotomy was performed using a 0.5-mm stainless steel drill bit (Meisinger), and a 4-mm diameter glass coverslip was placed over the exposed cortex and secured with dental cement. Postoperative analgesia was provided by buprenorphine (0.1-1 mg/kg) administered every 8 h.

### In vivo imaging

Chronic in vivo imaging of layer 1 OLs, myelin internodes, and OPCs was initiated 14-21 days after cranial window implantation in *Plp:GFP and Plp:GFP × Cspg4:dsRed* mice. Imaging was performed using an Olympus FV-RS microscope (Olympus, Japan) equipped with femtosecond pulsed Ti:Sapphire lasers (Insight and Mai Tai HP-DS, Spectra-Physics) and a resonant scanner with 16× line averaging. Laser power did not exceed 30 mW at the back focal plane.

For each imaging session, mice were anesthetized with 4-5% isoflurane for induction and maintained at 0.8-1.5% isoflurane during imaging. Physiological parameters, including anaesthesia depth, blood oxygen saturation, and heart rate, were continuously monitored using a MouseOx system (Starr Life Sciences) with a thigh sensor.

During the initial imaging session, 5-10 regions of interest (203 × 203 × 80 µm) were randomly selected across the somatosensory cortex and imaged at a spatial resolution of 0.4 µm (x/y) and 0.5 µm (z). In experimental regions, either one (single ablation) or three (multiple ablations) OLs were ablated by targeting the soma center using tornado-mode laser scanning (920 nm, 80-120 mW, 1 s). Ablated regions were re-imaged 2-3 days later to confirm OL death. All regions were subsequently imaged every 2-14 days for up to 91 days. Control regions were imaged over the same period without ablation. Regions exhibiting deteriorating imaging quality during longitudinal imaging were excluded from analysis; no evidence of photodamage was observed.

### Application of pharmacological agents

Systemic pharmacological treatment was initiated following laser-induced demyelination and continued for three weeks. Mice received daily subcutaneous injections of the CXCR4 antagonist AMD3100 (5 mg/kg; Merck A5602) diluted in 1× PBS. Injections were administered on the dorsal flank using a 30-gauge needle.

### Image processing and OL / myelin analysis

Post-processing of two-photon imaging data was performed using ImageJ/Fiji (version 2.14.0/1.54h; http://fiji.sc (Schneider et al., 2012)). Brightness and contrast were adjusted uniformly across images, and image stacks were aligned using the StackReg plugin.

Quantification of newly differentiated OLs following single or multiple ablations, as well as in control regions, was performed on in vivo recordings spanning 83-87 days. For spatial analyses in ablated regions, distances were calculated relative to the center of the ablated OL. In control regions, an OL in the center of the imaged area was defined as the reference point at day 0 (D0).

Myelin properties were analyzed for ablated OLs and for newly appearing OLs located more than 50 µm from the borders of the imaging volume. Three-dimensional reconstructions of lost and newly formed myelin internodes were generated within the chronic imaging volumes using the Simple Neurite Tracer plugin in Fiji. Internodes extending beyond the imaging volume were included in analyses of remyelination efficiency, total number, and total length of gained or lost internodes, but were excluded from internode-type classification and length-per-type analyses. The number and length of lost internodes were determined from the initial imaging time point (D0), whereas newly formed internodes were quantified at the final imaging session. Remyelination efficiency and internode type were determined relative to the D0 reference image.

### OPC dynamics and fate analysis

OPC dynamics were analyzed using in vivo recordings acquired at least twice per week over three-week intervals (D0-21 or D28-49 after imaging start) in both ablated and control regions. Maximum intensity projections (203 × 203 × 40 µm) were generated for each time point, assembled into image stacks, and aligned using StackReg prior to tracking. Individual OPCs were tracked manually over time using the Manual Tracking plugin in Fiji.

The distance traveled by each OPC was calculated from tracked coordinates according to

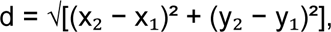

and OPC motility (µm/day) was determined by dividing the total distance traveled by the duration over which the cell could be reliably tracked. Only OPCs visible for at least two consecutive imaging sessions were included. Cells were tracked until two imaging sessions prior to death or differentiation, as indicated by morphological changes. Proliferating OPCs were tracked until two imaging sessions before and/or after cell division.

OPC fate was determined based on longitudinal in vivo morphology. OPCs were classified as proliferating when a single OPC soma observed at one imaging time point underwent division into two daughter cells at the subsequent imaging session. OPCs were classified as dying when an OPC soma exhibited acute swelling or fragmentation followed by complete disappearance, or direct loss of the soma accompanied by persistent cellular remnants at the subsequent imaging session. Cells migrating out of the imaging volume were not classified as dying.

### Statistics

All data are presented as mean ± S.E.M. Statistical analyses were performed using GraphPad Prism (version 10.6.1). Data distribution was assessed for normality, and appropriate parametric or nonparametric statistical tests were applied as indicated in the figure legends. The number of animals, imaging areas, cells, and myelin internodes analyzed for each experiment is reported in the corresponding figure legends. Investigators were blinded to age and experimental conditions during data analysis, where feasible. Statistical significance was defined as *P* < 0.05.

## Use of generative AI

We used large-language model (LLM)-based software (ChatGPT 5, Perplexity, Microsoft Copilot, and Grammarly) as search engines and for language editing. All results were checked and edited by the authors. No data analysis or figure material was generated using LLMs.

## Data availability

The datasets generated and analyzed during the current study are available from the corresponding authors on reasonable requests. Mouse lines can be requested under academic material transfer agreements from The Jackson Laboratories or from the laboratories that generated the lines, as indicated.

## Acknowledgements

We are grateful to D. Steinmetz and N. Budak for excellent animal husbandry and K. Kellermann for veterinary support. We thank Y. Hufnagel, M. Schetterer, and K. Wullimann, F. Beyer and S. Berger for outstanding technical, data management and administrative support. We also thank B. Zalc (ICM Paris) for his generous provision of *PLP:GFP(z)* mice (Spassky et al., 2001).

This project was supported by funding to N.S. from the Hertie Network of Excellence in Clinical Neuroscience (P1200019) and the Else Kröner-Fresenius-Stiftung (2022_EKEA.162). Additional support to N.S. was provided by the Hertie Foundation as part of the independent research group leader funding scheme. Intravital and high-resolution microscopy was supported via DFG instrumentation grants (INST 37-1170-1 FUGG – ID 467868227 and INST 95/1755-1 – FUGG, ID 518284373). Work in T.M.’s lab is supported by the Deutsche Forschungsgemeinschaft (TRR 274/1-2 2020, project B03 and Z01 – ID 408885537, Mi 694/9-1/2 A03 – ID 405358801, FOR Immunostroke, TRR 167/3 2025 – ID 259373024, and CRC 1744/1 2026 – ID 548585053). T.M. receives support from the Munich Center for Systems Neurology (SyNergy EXC 2145 – ID 390857198) and the German Center for Neurodegenerative Diseases. K.E. and S.A. are members of the Graduate School of Systemic Neurosciences (GSN), and L.S. was supported by the Elite Network Bavaria Master program Biomedical Neuroscience (BmN).

## Author contributions

T.M. and N.S. conceptualized the project and experiments. K.E. performed most of the investigation, including in vivo imaging and structural microscopy, and analyzed most of the data, with support from L.S. S.A. supported the management of mouse lines and in vivo imaging. N.S. and T.M. were responsible for funding acquisition; K.E., T.M., and N.S. wrote the paper with input from all authors.

## Extended Data figure legends

**Extended Data Figure 1.**
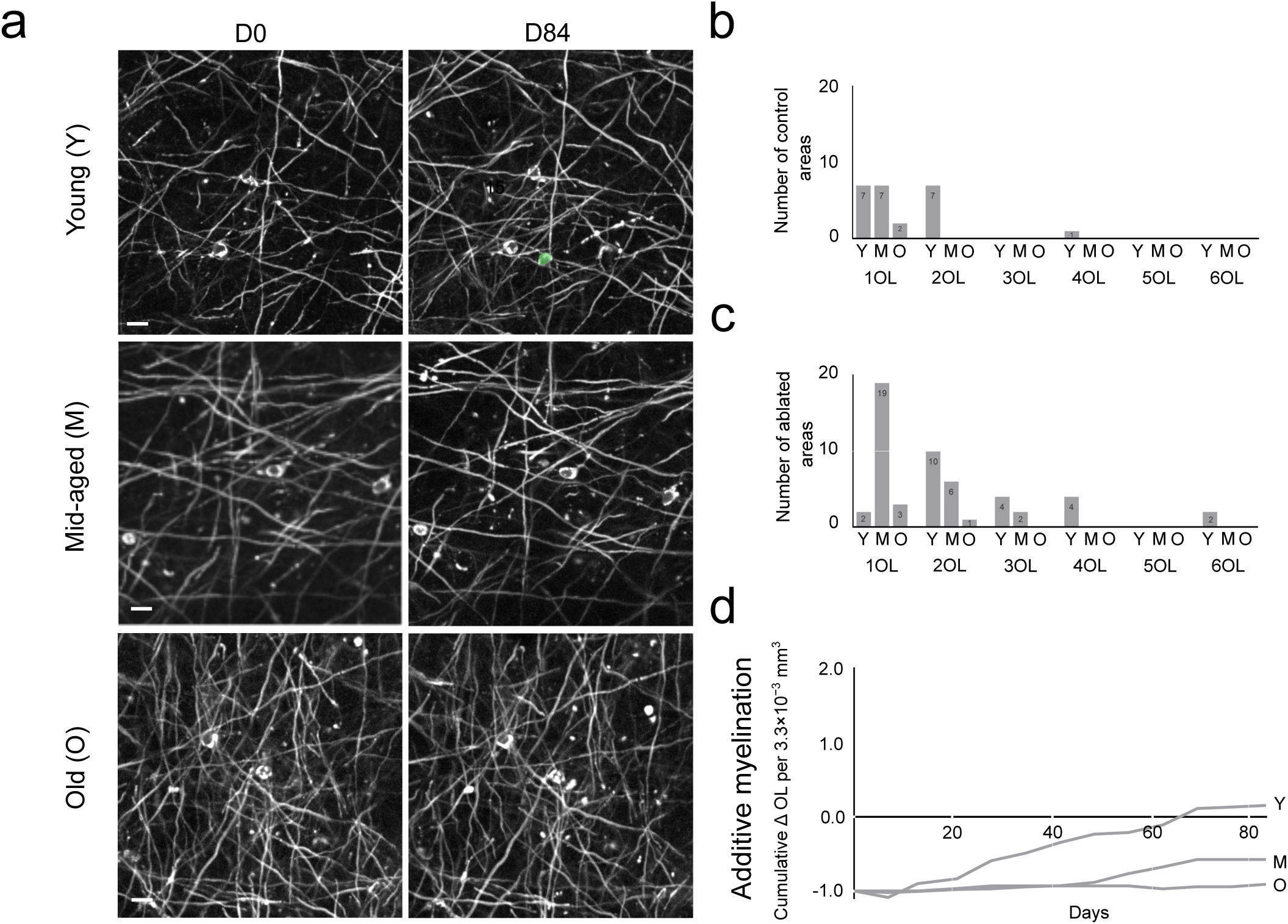
Distribution of newly differentiated oligodendrocytes in control and ablated cortex across age groups. **a,** In vivo imaging in young (top), mid-aged (middle), and old (bottom) mice. Left, imaging area at day 0. Right, the same region at later time points, with newly differentiated OLs shown in green where present. Only a subset of the full imaging area is shown. Maximum-intensity projections, 5 µm. **b,** Number of control areas (3.3×10⁻³ mm³) containing 1-6 newly differentiated OLs during the imaging window across age groups (Y; M; O). **c,** Number of ablated areas (3.3×10⁻³ mm³) containing 1-6 newly differentiated OLs during the imaging window across age groups (Y; M; O). **d,** Cumulative number of additional newly differentiated OLs per 3.3×10⁻³ mm³ over time across age groups. Values were calculated by subtracting the number of new OLs in control areas from those in ablated areas; the original control and ablated curves are shown in Fig. 1. Bars = mean ± SEM; n = young: 15 OL+ control areas and 22 OL+ ablated areas from 8 mice; mid aged: 7 OL+ control areas and 27 OL+ ablated areas from 18 mice; old: 3 OL+ control and 4 OL+ ablated areas from 10 mice. Scale bar, 10 µm (b). See Methods for details.

**Extended Data Figure 2.**
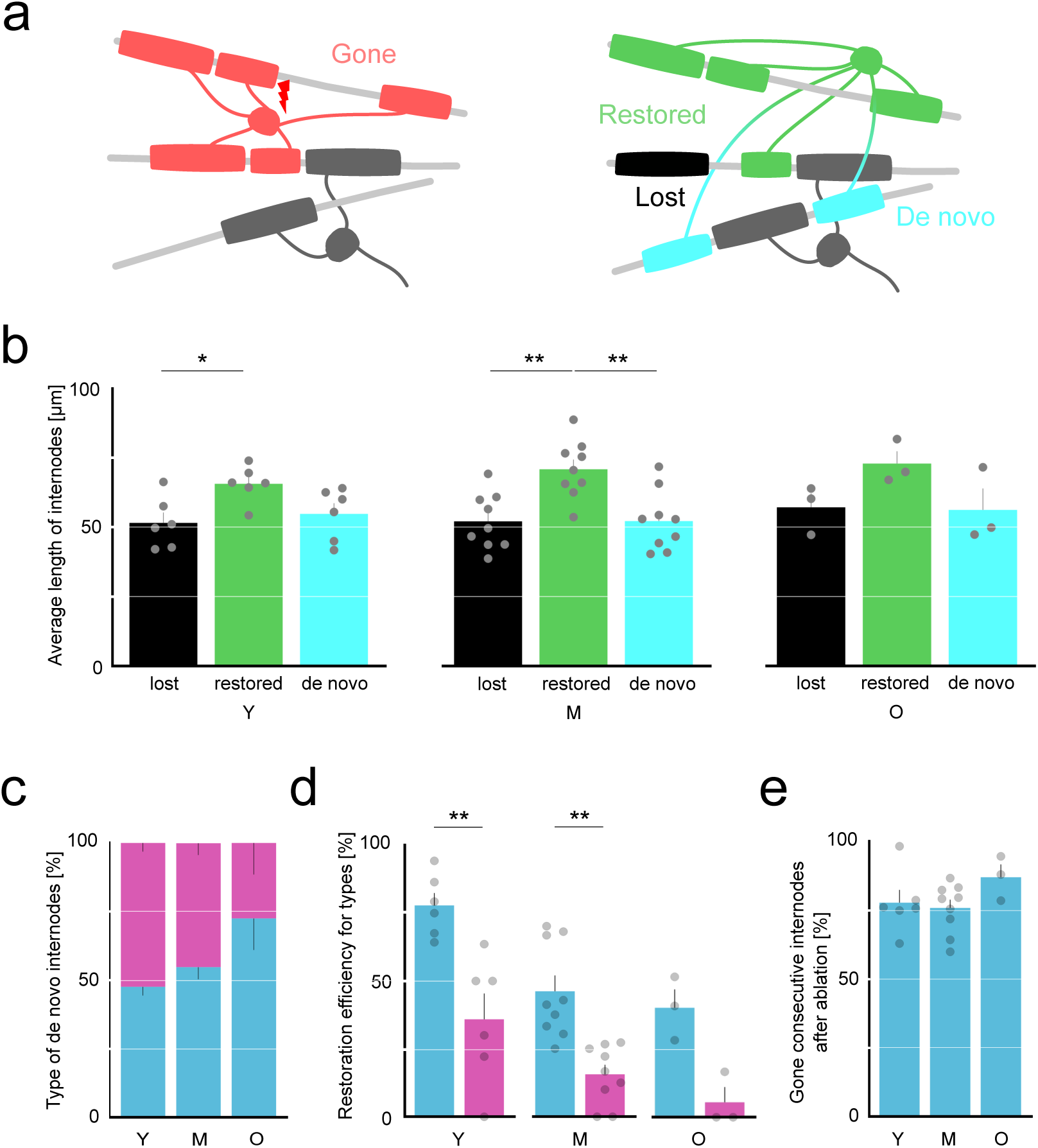
Age-dependent characteristics of internode loss and restoration. **a,** Schematic representation of demyelination (left, red), remyelination (green), and de novo myelination (cyan) by a newly generated oligodendrocyte (OL). Lost, non-restored myelin internodes are shown in black. Grey OLs and internodes indicate stable OLs; light grey lines represent axons. **b,** Average length of lost (black), restored (green), and de novo (cyan) myelin internodes after ablation across age groups. **c,** Fraction of de novo myelin internodes classified as consecutive (blue) or isolated (purple) in young, mid-aged, and old animals **d,** Fraction of consecutive (blue) and isolated (purple) myelin internodes that were restored. **e,** Fraction of demyelinated consecutive internodes after ablation. Bars = mean ± SEM; dots = individual imaging areas. n = young: 6 areas from 5 mice; mid-aged: 9 areas from 9 mice; old: 3 areas from 2 mice. Statistics: Standard (ordinary) one-way ANOVA with Tukey’s multiple comparisons test (b,e); unpaired two-tailed t test (d). Significance is indicated as P < 0.05 (*) and P < 0.01 (**); higher significance levels are collapsed into **. Exact P values: b (left), lost vs restored, P = 0.03; lost vs de novo, P = 0.79; restored vs de novo, P = 0.10; b (middle), lost vs restored, P = 0.002; lost vs de novo, P > 0.99; restored vs de novo, P = 0.002; b (right), lost vs restored, P = 0.22; lost vs de novo, P = 0.99; restored vs de novo, P = 0.19; d (left), consecutive vs iso (Y), P = 0.003; d (middle), consecutive vs iso (M), P = 0.002; d (right), consecutive vs iso (O), P = 0.25; e, Y vs M, P = 0.93; Y vs O, P = 0.39; M vs O, P = 0.23; See Methods for details.

**Extended Data Figure 3.**
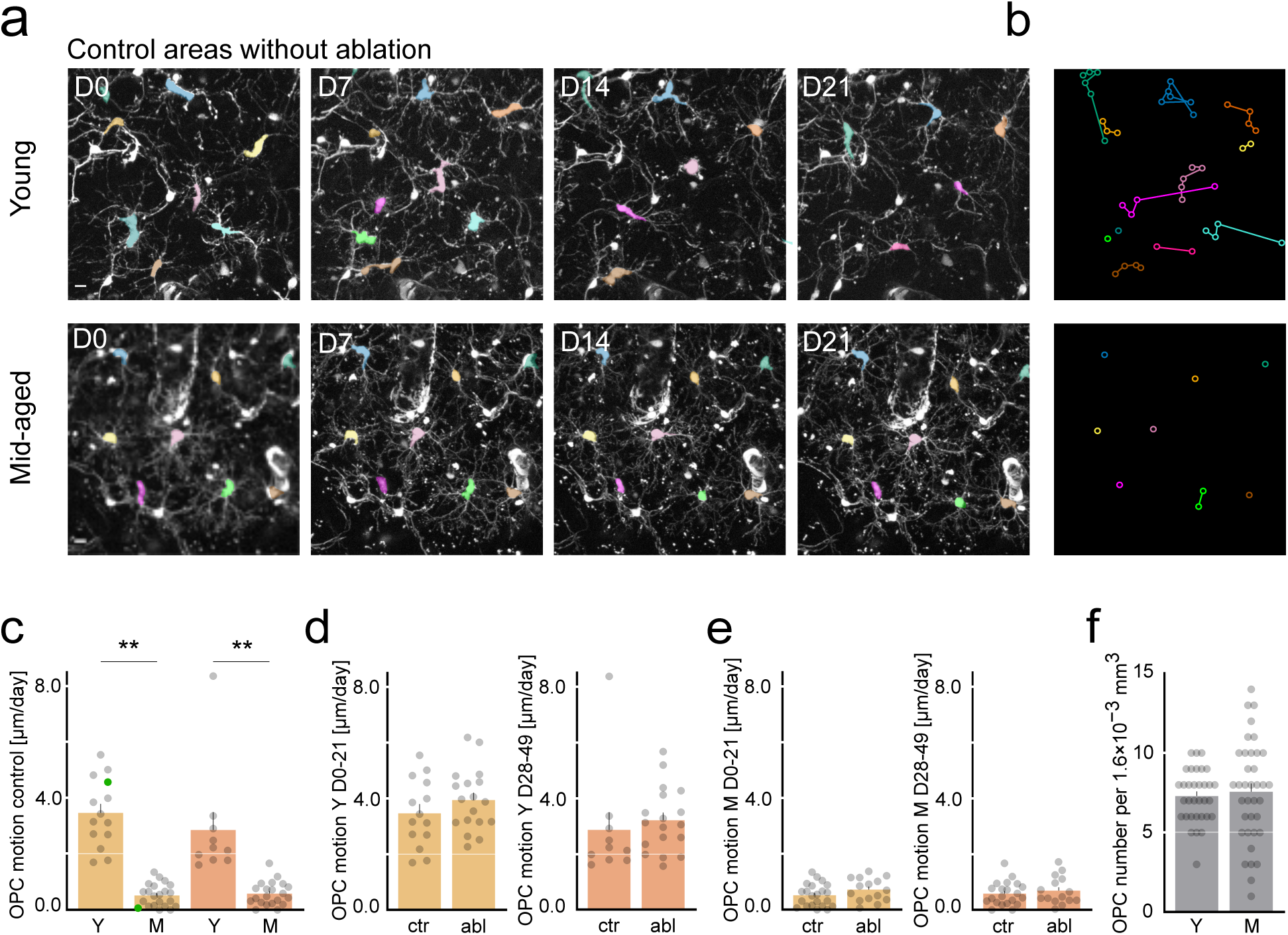
OPC motility in control and ablated cortex across age groups. **a,** In vivo longitudinal two-photon imaging of OPCs in control areas of young (top) and mid-aged (bottom) mice. OPC somata present for at least two consecutive time points were analyzed and pseudo-colored consistently across imaging sessions. Representative images are shown. Maximum-intensity projections, 40 µm. **b**, Trajectories of OPC soma movement derived from the representative images in **a**. Arrowheads indicate the direction of movement between consecutive time points while open circles indicate stable OPCs. **c**, Average daily OPC soma displacement in young compared to mid-aged control areas before (light orange) and during (dark orange) the time window corresponding to the de-/remyelination phase in the ablated areas shown in Fig. 3. **d**, Average daily OPC soma displacement in young mice comparing control and ablated areas before (light orange, left) and during (dark orange, right) the time window corresponding to the de-/remyelination phase. **e**, Average daily OPC soma displacement in mid-aged mice comparing control and ablated areas before (light orange, left) and during (dark orange, right) the time window corresponding to the de-/remyelination phase. **f**, Average OPC number in young compared to mid-aged ablated areas at day 0. Bars = mean ± SEM; dots = individual imaging areas. Colored dots indicate values corresponding to the representative images shown in a. n = young: 18 ablated areas from 5 mice; mid-aged: 15 ablated areas from 6 mice. Statistics: one-way ANOVA with Tukey’s multiple comparisons test (c), two-tailed t-test (d-f) Significance is indicated as P < 0.05 (*) and P < 0.01 (**); higher significance levels are collapsed into **. Exact P values: c, Y(before) vs M(before), P < 0.0001, Y(before) vs Y(during), P = 0.46, Y(during) vs M(during) P < 0.0001, M(before) vs M(during) P > 0.99; d (left), control vs ablated P = 0.26, d (right), control vs ablated P = 0.56, e (left), control vs ablated P = 0.15, e (right), control vs ablated P = 0.49, f, Y vs M P = 0.67. All post hoc tests were performed, but only comparisons relevant to the experimental question are shown. Scale bar, 10 µm (a). See Methods for details.

**Extended Data Figure 4.**
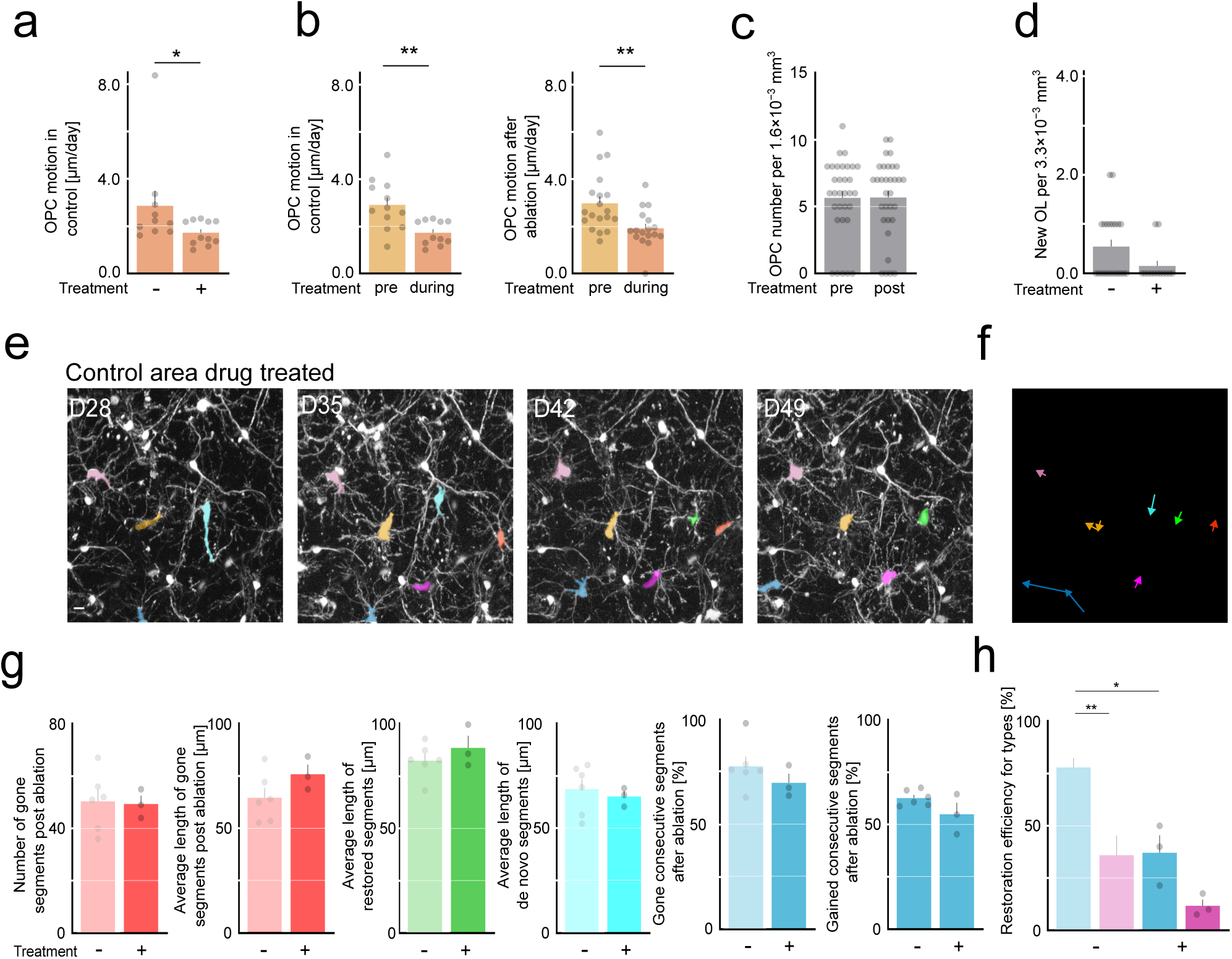
OPC dynamics and myelin internodes properties in CXCR4-antagonist treated and untreated animals. **a,** Average daily OPC soma displacement during the de-/remyelination phase in control areas. **b**, Average daily OPC soma displacement in control areas (left) before (light orange) and during (dark orange) the time window corresponding to the de-/remyelination phase in ablated areas, and in ablated areas (right) before (light orange) and during (dark orange) the de-/remyelination phase. **c,** Average OPC number in young CXCR4-antagonist treated animals at day 28 (pre) and day 49 (post). Control and ablated areas were merged. **d**, Density of newly differentiated OLs per 3.3×10⁻³ mm³ during the treatment window in CXCR4-antagonist treated compared to untreated animals in control regions. **e**, In vivo longitudinal two-photon imaging of OPCs in a control area of a young CXCR4-antagonist treated mouse. OPC somata present for at least two consecutive time points were analyzed and pseudo-colored consistently across imaging sessions. Representative images are shown. Maximum-intensity projections, 40 µm. **f**, Trajectories of OPC soma movement derived from the representative images in e. Arrowheads indicate the direction of movement between consecutive time points. **g**, from left to right: number of demyelinated myelin internodes, average length of demyelinated myelin internodes, average length of restored myelin internodes, average length of de novo myelin internodes, fraction of demyelinated consecutive internodes, and fraction of restored consecutive internodes after ablation in young untreated (Graphs 1-4 replots from Fig. 2 and Extended Data Fig. 2) versus CXCR4-antagonist treated animals. **h**, Fraction of restored myelin internodes classified as consecutive (blue) or isolated (purple) in untreated (replot from Extended Data Fig. 2d) versus CXCR4-antagonist treated young animals. Bars = mean ± SEM; dots = individual imaging areas. Replotted values from Figs. 2 and 3 are shown without individual data points and greyed out. n (OPC dynamics): control untreated (during) = 10 areas from 5 mice; control treated (during) = 11 areas from 5 mice; control treated (pre) = 12 areas from 6 mice. n (OPC number) = 32 areas from 12 mice. n (myelin analysis, g-h) = treated: 3 areas from 3 mice. Statistics: two-tailed Mann-Whitney U test (a,c,, d, g, graphs 5-6); paired two-tailed t test (b); unpaired two-tailed t test (g, graphs 1-4); Welch’s one-way ANOVA followed by Dunnett’s T3 multiple-comparisons test (h). Significance is indicated as P < 0.05 (*) and P < 0.01 (**); higher significance levels are collapsed into **. Exact P values: a, – vs +, P = 0.026; b, – vs +, P = 0.93; c (left), pre vs during, P = 0.002; c (right), pre vs during, P = 0.002; d, P = 0.053; g (from right to left), – vs +, P = 0.89; – vs +, P = 0.18; – vs +, P = 0.36; – vs +, P = 0.64; – vs +, P = 0.55; – vs +, P = 0.26; h, consecutive (untreated) vs iso (untreated), P = 0.003; consecutive (untreated) vs consecutive (treated), P = 0.017; consecutive (treated) vs iso (treated), P = 0.28; iso (untreated) vs iso (treated), P = 0.21. All post hoc tests were performed, but only comparisons relevant to the experimental question are shown. Scale bar, 10 µm (**e**). See Methods for details.

**Extended Data Figure 5.**
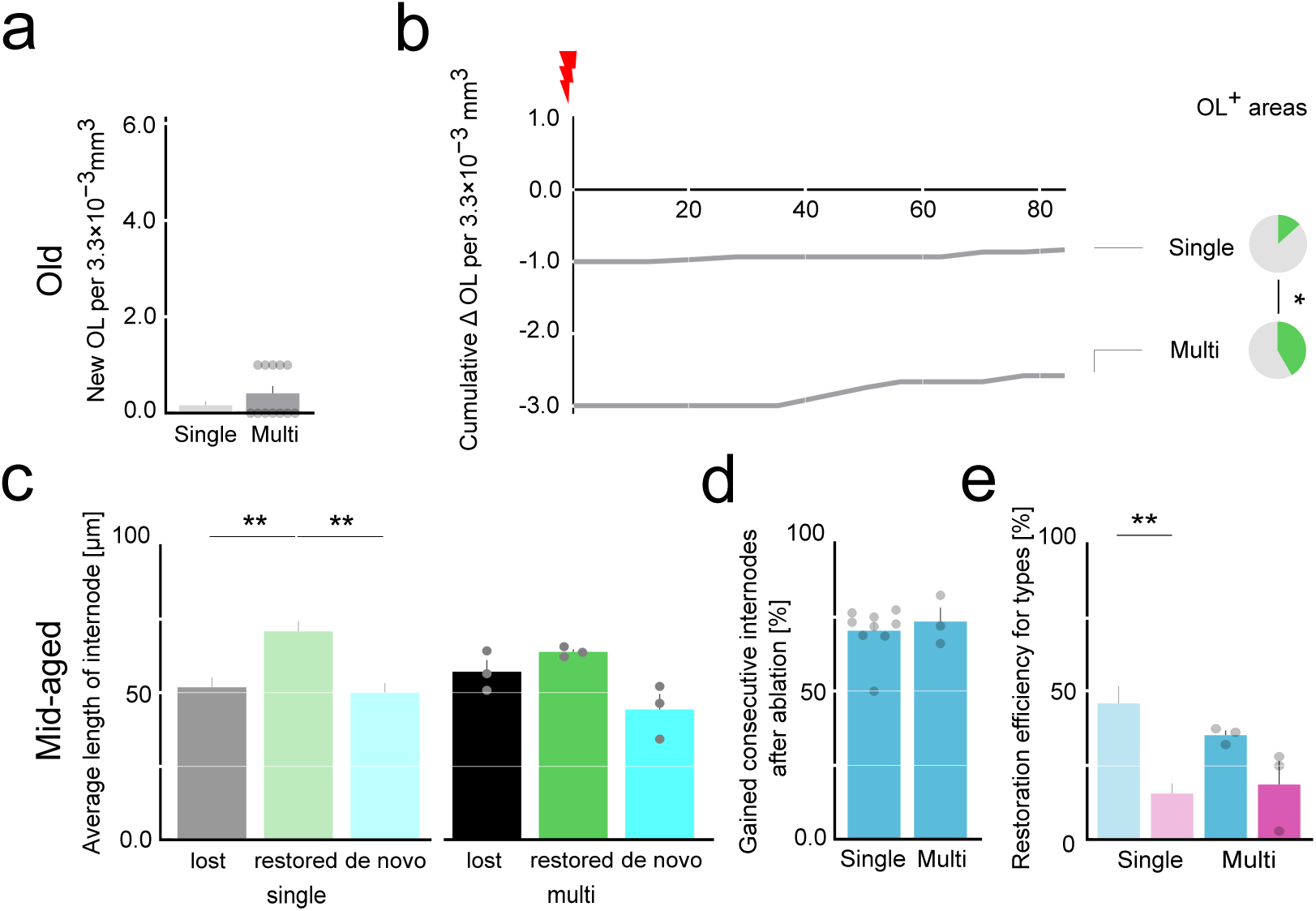
Oligodendrocyte generation and myelin internodes characteristics after single- and multiple-cell ablation across age groups. **a,** Number of newly differentiated oligodendrocytes (OLs) per 3.3×10⁻³ mm³ in regions following single-cell and multiple-cell laser ablation in old animals. **b,** Cumulative number of newly differentiated OLs per 3.3×10⁻³ mm³ over time in regions following single-cell and multiple-cell laser ablation in old animals. Pie charts show the fraction of “OL+ areas” (areas containing at least one newly differentiated OL) after single-cell ablation (top, replot from Fig. 1g) and multiple-cell ablation (bottom). **c,** Average length of lost (black), restored (green), and de novo (cyan) myelin internodes after ablation, comparing mid-aged single- (replot from Extended Data Fig. 2b) and multiple-ablated regions. **d,** Fraction of gained consecutive myelin internodes in mid-aged single- and multiple-ablated regions. **e,** Fraction of restored myelin internodes classified as consecutive (blue) or isolated (purple) in mid-aged single- (replot from Extended Data Fig. 2d) and multiple-ablated regions. Bars = mean ± SEM; dots = individual imaging areas. Replotted values from Figs. 1, 2 and Extended Data Fig 2 are shown without individual data points and greyed out. n (OL): multi old = 12 areas from 5 animals (a,b); n (myelin mid-aged multi): 3 areas from 3 animals (d-e). Statistics: two-tailed Mann-Whitney U test (a); two-sided chi-square test (b, pie charts); Welch’s one-way ANOVA followed by Dunnett’s T3 multiple-comparisons test (c, e); Welch’s two-tailed t-test (d). Significance is indicated as P < 0.05 (*) and P < 0.01 (**); higher significance levels are collapsed into **. Exact P values: a, single vs multi, P = 0.09; b, single vs multi, P = 0.043; c, lost (single) vs restored (single), P = 0.002; lost (single) vs de novo (single), P = 0.998; restored (single) vs de novo (single), P = 0.0008; lost (multi) vs restored (multi), P = 0.95; lost (multi) vs de novo (multi), P = 0.56; restored (multi) vs de novo (multi), P = 0.15; lost (single) vs lost (multi), P = 0.96; restored (single) vs restored (multi), P = 0.87; de novo (single) vs de novo (multi), P = 0.94; d, single vs multi, P = 0.62; e, consecutive (single) vs iso (single), P = 0.0007; consecutive (multi) vs iso (multi), P = 0.46; consecutive (single) vs consecutive (multi), P = 0.65; iso (single) vs iso (multi), P = 0.98. All post hoc tests were performed, but only comparisons relevant to the experimental question are shown. See Methods for details.

## Notes

### Competing Interest Statement

The authors have declared no competing interest.

